# Evidence from Formal Logical Reasoning Reveals that the Language of Thought is not Natural Language

**DOI:** 10.1101/2025.07.26.666979

**Authors:** Hope Kean, Alexander Fung, Paris Jaggers, Jason Chen, Joshua S. Rule, Yael Benn, Joshua B. Tenenbaum, Steven T. Piantadosi, Rosemary A. Varley, Evelina Fedorenko

## Abstract

Humans are endowed with a powerful capacity for inductive and deductive logical thought: we easily form generalizations based on a few examples and draw conclusions from known premises. Humans also arguably have the most sophisticated communication system in the animal kingdom: natural language allows us to express complex and structured meanings. Some have therefore argued for a tight relationship between complex thought and language, postulating that reasoning, including logical reasoning, relies on linguistic representations. We systematically investigated the relationship between logical reasoning and language using two complementary approaches. First, we used non-invasive brain imaging (fMRI) to examine neural activity as healthy adults engaged in logical reasoning tasks. And second, we behaviorally evaluated logical abilities in individuals with extensive lesions to the language brain areas and consequent severe linguistic impairment. Our findings reveal that the language brain network is not engaged during logical reasoning, and patients with severe aphasia exhibit intact performance on logic tasks. Instead, inductive reasoning recruits the domain-general multiple demand network implicated broadly in goal-directed behaviors, whereas deductive reasoning draws on brain regions that are distinct from both the language and the multiple demand networks. Together, these results indicate that linguistic representations are neither utilized nor required for inductive or deductive logical reasoning.

**Significance:** Which cognitive mechanisms allow humans to reason logically, to understand whether a conclusion follows from the premises? Are they the same ones that allow the assembly of words into structured representations? Scholars have debated for millennia whether logical reasoning is inextricably tied to natural language, or instead relies on a distinct “language of thought” (LOT). Using fMRI in healthy adults and evaluating logical ability in individuals with severe aphasia, we find that distinct neural systems support language processing vs. logical (inductive and deductive) reasoning. These results suggest that, at least in mature brains, language processing does not underpin logical inference, perhaps due to the distinct representational format of the logical LOT.

## Introduction

In 350 BC, Aristotle wrote that “writing and speech are not the same for all people, but mental acts themselves, of which words signify, are the same for all people” (1). This reflection highlights a distinction between linguistic symbols and the underlying cognitive structures they aim to express. But over the years, many philosophers, linguists, and cognitive scientists have argued that natural language is, in fact, the medium of complex thought (e.g. (2–18)); for counterarguments, see (19, 20). We here challenge this view using evidence from human neuroscience.

A key argument for language underlying thinking has to do with similarities between them. For example, one prominent proposal about thought—the “language of thought” (LOT) hypothesis (21)—has emphasized the compositional and hierarchical nature of thoughts. According to this hypothesis, thoughts are composed of smaller atomic pieces in a structured format with hierarchical relations among the component elements. A parallel to natural language is easy to draw: sentences are built out of words, which are related to one another hierarchically. But a parallel can also be drawn to computer programs, which are built out of a small collection of primitive operations, or to melodies, which are composed of hierarchically-related notes and chords. So, this similarity alone is insufficient to conclude that we think in language. Furthermore, although many thoughts can be cast into natural language (which is what makes language an effective communication system), we can also express ideas—including abstract ones—using non-linguistic symbols, mathematical expressions, visual schematics or diagrams, and so on. Therefore, a degree of isomorphism between certain thoughts and linguistic expressions need not entail that linguistic representations are used for thinking, any more than the possibility of expressing certain ideas using visual images entails that our thinking is visual in nature.

Indeed, empirical data have been accumulating that suggest that linguistic representations are neither utilized nor necessary for thinking. Some patients with aphasia appear capable of diverse forms of thought, as evidenced by their intact performance on tasks requiring mathematical reasoning (22), causal reasoning (23), and Theory of Mind (24, 25); (see (26) and (27) for reviews). Converging evidence for the separability of language and reasoning comes from brain imaging studies. The brain areas that support language comprehension and production (28–30) are strongly sensitive to linguistic syntactic structure (31–37) but are not engaged during non-linguistic cognitively demanding tasks, including those that require representing and manipulating structured representations (see (38) for a review). These tasks span solving arithmetic and algebraic problems (39–41), understanding computer code (42, 43), social reasoning (44–47), and performing demanding executive function tasks (39, 48, 49), which require skills linked to fluid intelligence (50, 51).

The separability of language and thought has been questioned by some researchers based on a) correlations between the degree of linguistic impairment and impairment on cognitive tasks in aphasia (e.g., (52–55)) and b) evidence from verbal interference dual-task paradigms (for a review, see (56)). We believe that these findings have straightforward alternative explanations and thus do not present a challenge for the claim that language and thought rely on distinct resources (see <u>Discussion</u>). However, there is one domain of thought whose relationship with language has only received limited attention in the prior literature, despite it being a hallmark domain where the role of language has been emphasized (11, 57). This domain is *abstract logical reasoning*.

One prior fMRI study examined the relationship between language and deductive reasoning by comparing neural responses during a logical inference task (judging the validity of a conclusion given a premise) versus a linguistic inference task (judging the validity of thematic roles across two sentences) relative to lower-level control conditions (judging expression well-formedness) (58). A partial dissociation was observed: although both the logic and the language contrast elicited left-lateralized responses in the frontal, temporal, and parietal cortex, some brain areas—in the vicinity of areas traditionally implicated in language processing (59–62, 38)—showed a stronger response to linguistic inference, whereas other areas responded more strongly during logical inference (see (63) for concordant evidence from TMS). However, recent evidence suggests that the kind of linguistic paradigm used by Monti et al. (58) engages both language-processing areas but also brain areas sensitive to general cognitive effort (e.g., (64)), which makes previously observed dissociations more difficult to interpret. Another past study examined logical and mathematical reasoning and argued that neither engaged the language areas (57), but that study did not include a language task, which would be necessary to make such a claim. Thus, these prior findings are intriguing, but have not conclusively answered the question of whether logical reasoning relies on linguistic processing mechanisms.

Two other bodies of work speak to the logic-language relationship. The first comes from developmental psychology and suggests that the timelines of linguistic and logic development diverge. Inductive reasoning appears to come online very early (e.g., (65, 66)). The representations are argued to be sensorimotor, not linguistic, in nature, and the main debate concerns their domain specificity (e.g., (67, 68)) vs. generality (e.g., (69, 70)) (e.g., do infants have distinct representations for reasoning about the physical world vs. about social agents?). For deductive reasoning, the picture is less clear: some have argued that preverbal infants can already reason disjunctively (71), which implies a separation of logical and linguistic ability. However, others have reported failures in simple disjunctive inferences as late as age 2.5 years (72), and even later for more complex inferences (73). These late failures may suggest that a certain level of linguistic competence is required for the development of these capacities, although they may also reflect the slow development of executive abilities (74–76), which may be needed to deal with the task demands in complex paradigms.

The second body of work comes from artificial intelligence, where recent advances have provided additional motivation for investigating the relationship between language and logic. In particular, neural network language models (77–79) not only achieve human-level performance on diverse language tasks (80–82), but also exhibit impressive successes on certain reasoning tasks (e.g., ARC: (83); BIG-Bench: (84); LogiQA: (85); MATH: (86); WildBench: (87); Templates: (88)). These findings beg the question of whether linguistic competence inevitably leads to logical ability, although many have pointed out that reasoning in large language models lacks robustness and generalizability (e.g., (89–94)).

Thus, the relationship between language and logic—and more specifically, whether abstract logical reasoning relies on linguistic representations in humans—remains unclear. To systematically examine the role of language in human logical reasoning, we adopted a two-pronged approach. We first used fMRI in healthy adults to test whether the language network (38) responds to logical reasoning demands (Study 1). Next, we used behavioral experiments in individuals with severe aphasia resulting from extensive damage to left peri-Sylvian cortex, to test whether linguistic ability is necessary for logical reasoning (Study 2). Note that our research does not speak to the potential role of language in the acquisition of logical concepts or in learning to reason during childhood (we briefly touch on this point in the <u>Discussion</u>). To foreshadow our findings, we find no evidence for the idea that abstract logical reasoning draws on linguistic resources.

## Results

In each study, we examined two paradigmatic forms of logical reasoning: inductive and deductive reasoning. The ***inductive reasoning*** paradigm (**Figure 1**, left) was adapted from Rule et al. (95) (for related paradigms, see (96–100)). Participants were presented with an input number list and an output list (e.g., `[5, 7] → [7, 5, 7]`) and asked to infer the rule that governs the input-to-output transformation. They could then test their hypothesis on a new input list, and so on, until they guess the correct rule. The rules involve a combination of mathematical operations (e.g., *f* (*l*) = [*x* + 2 ∣ *x* ∈ *l*]), “add 2 to every number”), list operations (e.g., *f* (*l*) = [*xi* ∣ *xi* ∈ *l*, ∀*j* < *i*, *xi* ≠ *xj*], “leave only unique, non-repeated elements”), and structural operations (e.g., *f* (*l*) = *l*[3:]*^l*[0:3], “rotate the list by three elements”), each of which can be written as a short computer program. The fMRI version also included a control condition where participants were told the rule and asked to apply it to new input lists.

**Figure 1.**
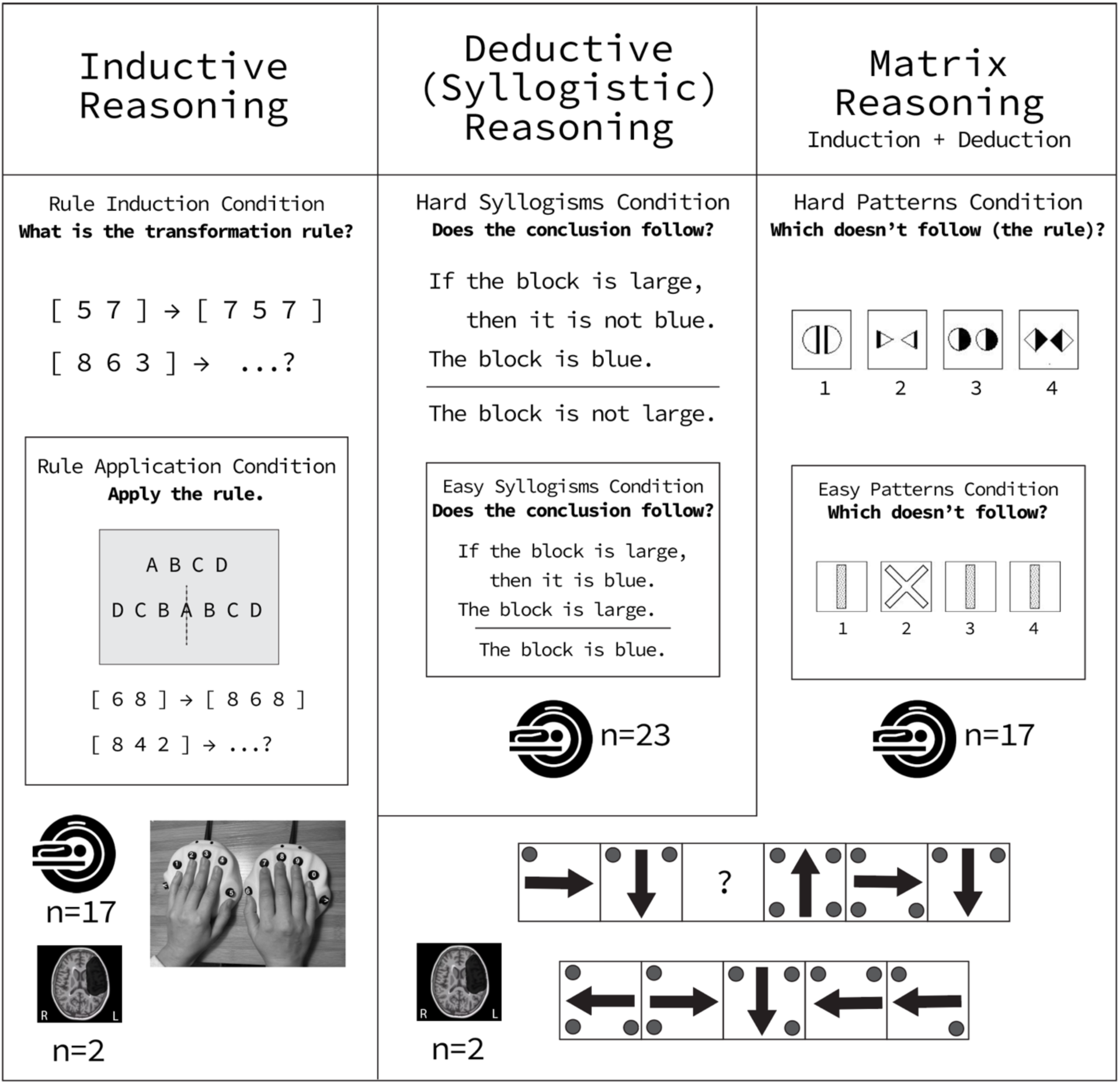
Logical reasoning paradigms. In the ***inductive reasoning*** task (left), participants are shown an input list of 1-5 single-digit numbers and an output list of 0-5 single-digit numbers and asked to guess the rule that transformed the input list into the output list (see <u>Methods</u> for details). In the fMRI version, after the eighth problem, participants were told—via a schematic— the correct rule and were asked to apply this rule to two more input lists. In the ***deductive syllogistic reasoning*** task (middle), participants are presented with classic three-sentence syllogisms and asked to judge the validity of the third sentence (the conclusion) given the first two sentences (the premises). Trials vary in difficulty between harder deduction (Modus Tollens) and easier deduction (Modus Ponens). In the ***matrix reasoning*** task (right), participants are presented with images of geometrical shapes and asked to figure out the rule. In the fMRI version, they are asked to decide which image does not fit with the others, and in the behavioral version, they have to identify the missing image (see <u>Methods</u> for details).

For ***deductive reasoning***, we used two paradigms: syllogistic reasoning and matrix reasoning. The first paradigm was introduced in Coetzee & Monti (101) and was only used here for the fMRI component of the study (**Figure 1**, center). Participants were presented with a classic syllogism consisting of two premises and a conclusion (e.g., i. If A is B, then A is also C. ii. A is not C. iii. Therefore, A is not B) and were asked to judge the validity of the conclusion. Half of the syllogisms used real words (e.g., “If the block is large…”), and the other half used nonwords (e.g., “If the tep is ag…”). Although deductive reasoning and language processing are required for all problems, the problems critically vary in the difficulty of the deduction. The easier problems, known as Modus Ponens, had the following forms: “If the block is large, then it is not yellow. The block is large. So, the block is not yellow.” (the conclusion follows from the premises), or “If the block is large, then is it not yellow. The block is large. So, the block is yellow.” (the conclusion does not follow from the premises). The more complex problems, known as Modus Tollens, had the following forms: “If the block is large, then it is not yellow. The block is yellow. So, the block is not large.” (the conclusion follows from the premises), or “If the block is large then it is not yellow. The block is not yellow. So, the ball is large.” (the conclusion does not follow from the premises).

The syllogistic paradigm is a classic paradigm in logical reasoning research (102–106) and the contrast between the Modus Tollens versus Modus Ponens problems elegantly isolates deductive demands. However, this task is not suitable for patients with aphasia because it uses verbal stimuli. As a result, for the second paradigm, we chose a non-verbal task, which could also be performed by the patients: matrix reasoning (107) (for prior fMRI use, see (108)). Participants were presented with a matrix of four geometric patterns and asked to find an outlier (**Figure 1**, right). In the control condition, the task is similarly structured but does not require much logical reasoning and can be performed using visual pattern recognition. Patients with aphasia performed a similar version (109), except that a matrix of geometric patterns was missing a pattern, and participants had to decide which option from several possible answers completes the matrix (**Figure 1**, bottom right panel). This type of paradigm, commonly used in assessing fluid reasoning capacity across diverse populations (110–119), taxes both deductive and inductive reasoning (120, 121). Inductive reasoning is needed to make guesses about the underlying rules, and deductive reasoning—to evaluate logical constraints and eliminate incorrect options. Thus, although not as pure as the syllogistic reasoning task, this paradigm is still well-suited to test the necessity of language for deductive reasoning.

If linguistic representations underlie logical inference, then a) the language brain areas should be sensitive to inductive demands, deductive demands, or both, as measured with fMRI; and b) patients with profound aphasia should show impairments on these tasks.

### The language network is not engaged during inductive, deductive, or matrix reasoning

The language-processing brain areas show little or no response to the inductive, deductive (syllogistic) and matrix reasoning contrasts. As expected based on much prior work (e.g., see (38) for a review), these areas showed a robust response to language processing, evidenced by a stronger response during the sentence compared to the nonword-list condition estimated in independent data (*p*<0.001 at the network level; **Figure 2B**, **Table 1**). Note that although a particular localizer contrast is used here to define the language areas, this contrast is highly generalizable across presentation modalities, languages, stimuli, and tasks (e.g., (122, 123, 48, 124)), and this network can also be defined without a contrast, using patterns of functional connectivity (e.g., (125–127)). Moreover, these language-responsive areas have been shown to be robustly sensitive to linguistic processing demands in both comprehension and production, and across controlled and naturalistic paradigms (review: (38)), and this ubiquity of engagement across diverse language tasks aligns with the fact that damage to these areas leads to language deficits (60, 128).

**Figure 2.**
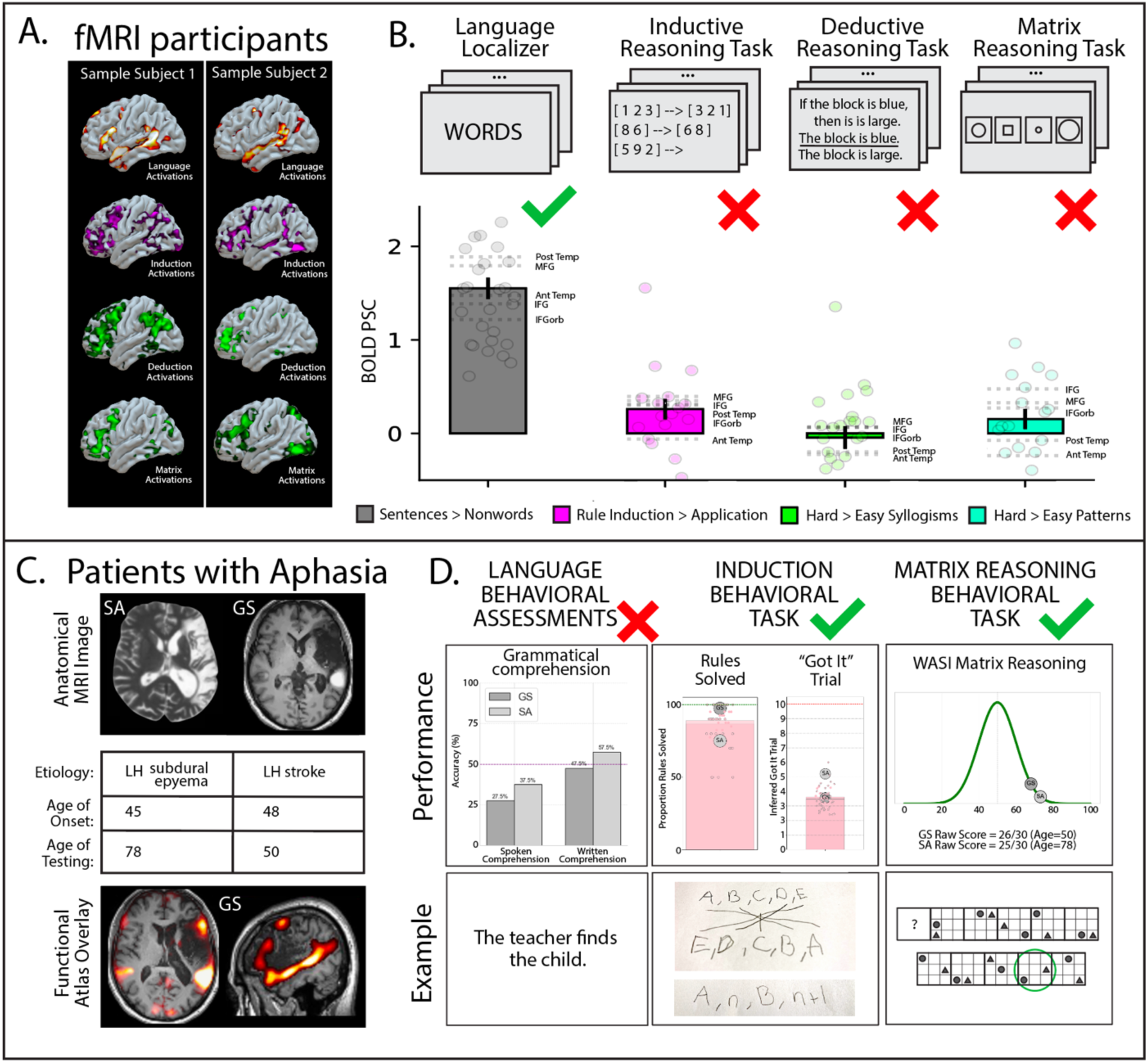
The language network does not support logical reasoning. **A.** Activation maps for the language contrast (1^st^ row, yellow) and three logical reasoning contrasts (2^nd^-4^th^ rows, purple and green) in two individual sample participants. **B.** Responses in the language network (individually defined in each participant; see <u>Methods</u>), in percent BOLD signal change, to the language contrast (sentences > nonwords; estimated in left-out data; grey), the induction contrast (rule induction > rule application; magenta), the deduction contrast (hard syllogisms > easy syllogisms; green), and the matrix contrast (hard patterns > easy patterns; teal). The bars show the across-network averages and the error bars correspond to the standard error of the mean by participant; the light dots correspond to individual participants’ responses; the means for each of the 5 language fROIs (IFGorb, IFG, MFG, AntTemp, and PostTemp) are shown with horizontal dashed lines. **C.** The anatomical scans of the two patients with severe aphasia (S.A. and G.S.), brief info on the patients, and a projection of the probabilistic atlas for the language network (from (266)) into the MRI image of G.S. **D.** Behavioral performance of the patients with aphasia and—for the critical tasks—control participants. The first column shows performance on the language evaluation tasks (spoken and written sentence comprehension; (267)) for S.A. (light grey bars) and G.S. (dark grey bars). Both participants show at or below chance performance. In the bottom row, we show a sample item. (See <u>Methods</u> for additional details of the linguistic evaluation of the patients.) The second and third columns show performance on the induction and matrix reasoning tasks (S.A.: light grey dots; G.S.: dark grey dots). For the induction task, the control data are shown with a pink bar. In the bottom row, we show two sample rule representations generated by G.S., where he illustrates the “reverse the numbers” rule (top) and the “insert index after number” (bottom). For the matrix reasoning task, the control data are shown with the normal age-matched distribution, and in the bottom row, we show a sample item.

**Table 1.**
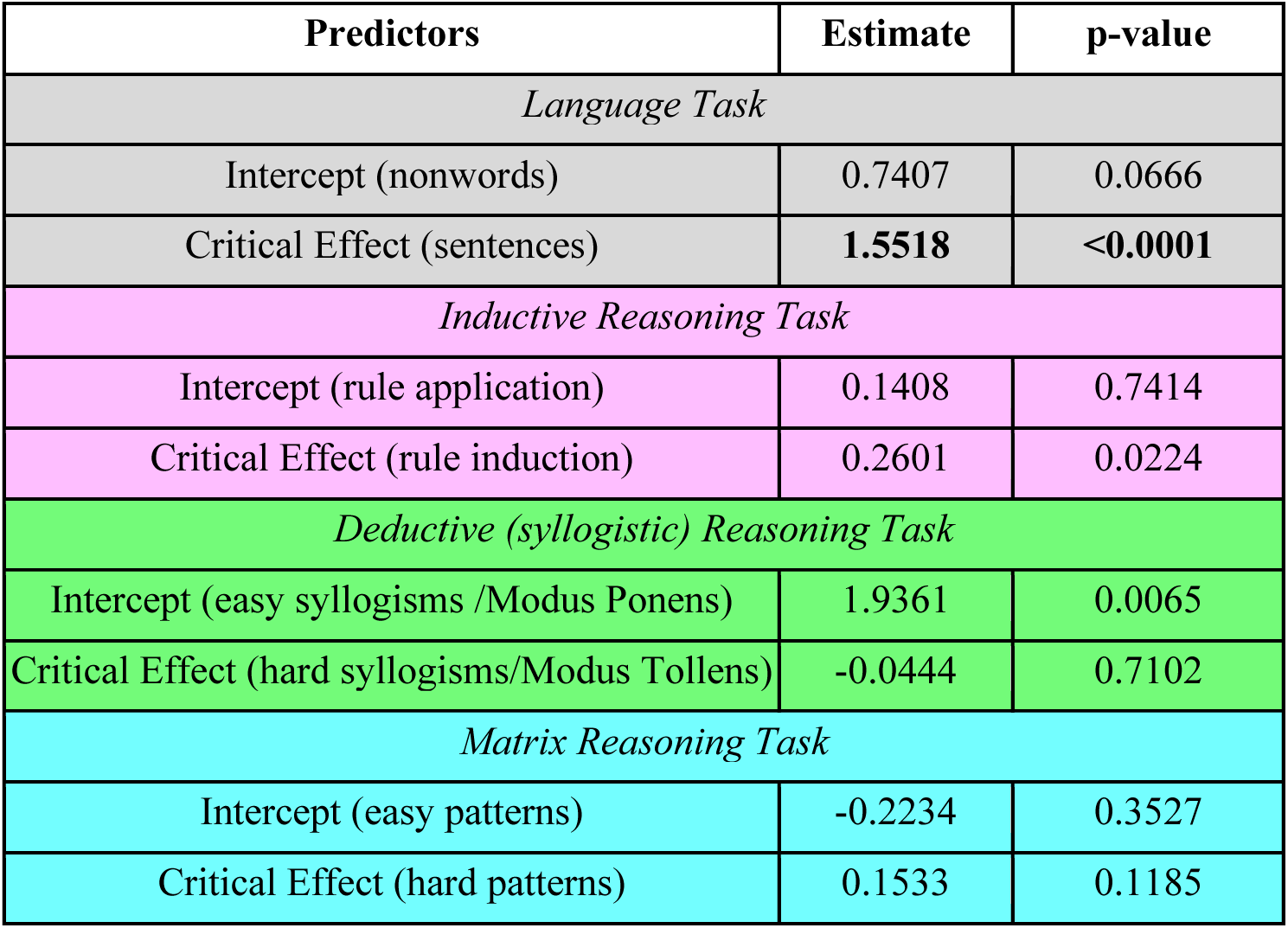
The responses of the language network to the language and logical reasoning contrasts. The results from the linear mixed-effects models (see <u>Methods</u>) at the network level (see **SI** Table 1 for the results broken down by fROI).

Critically, the language areas showed little or no sensitivity to the logic contrasts. The inductive contrast did elicit a reliable response at the network level (*p*<0.05), but the effect was small, with the language contrast being over four times stronger, and the critical induction condition eliciting a response at or below the level of the control condition of the language task (reading nonwords) (**SI Figure 1B**). For the remaining tasks, neither the deductive (syllogistic) nor the matrix reasoning contrast elicited a significant response in the language areas (*p*s>0.1 at the network level; **Figure 2A-B**, **Table 1**; see **SI Figure 1A** and **SI Table 1** for evidence that the results are similar for the five language areas examined separately), despite significant behavioral difficulty effects in both accuracies and reaction times for both tasks (**SI Figure 4**). For the syllogistic task, the responses relative to the low-level (fixation) baseline were strong to both the Modus Ponens and the Modus Tollens conditions, and comparable to the level of response to the sentences in the language localizer, which uses similar-length sentences but with no logic-related content (**SI Figure 1B**). This result is expected given that the processing of syllogisms requires lexical and syntactic processing). The critical result of no sensitivity to deductive reasoning difficulty generalized across words and nonwords (**SI Figure 5**). Interestingly, the conditions that use real words did not elicit a stronger response than the conditions that use nonwords in the language areas. The nonword conditions (“If the tep is ag…”) are similar to the so-called Jabberwocky sentences (129), which in past fMRI studies have been shown to elicit a lower response than real sentences in the language areas (e.g., (37, 122, 130, 131)). We speculate that this effect is absent here because the words’ meanings are irrelevant to the task, and, as a result, participants treat both words and nonwords as variables and parse the sentences in similar ways, extracting basic relational structures, which can then be ‘passed’ to the brain system that supports this type of reasoning (see <u>Inductive and deductive reasoning elicit strong responses outside the language network</u>).

### Patients with profound aphasia can nevertheless reason logically

Two individuals with profound aphasia, S.A. and G.S., showed preserved logical reasoning abilities. Both individuals have sustained extensive damage to left peri-Sylvian cortex (**Figure 2C**) and, consequently, exhibited severe linguistic impairments (**Figure 2D**, **SI Table 3**). In particular, critical to the hypothesis that linguistic syntactic structures mediate logical reasoning, these patients showed severe grammatical impairments in both comprehension and production, with at or near-chance performance on multiple syntactic assessments. For example, both exhibited marked impairment in assigning thematic roles in understanding reversible sentences, such as *The diver splashed the dolphin*. Nevertheless, despite their profound aphasia, both patients showed typical-like performance on the inductive and matrix reasoning tasks. In the inductive reasoning task (**Figure 1**, left), they solved the vast majority of the rules presented to them (S.A.: 19/25; G.S.: 39/40). This performance level is comparable to a normative sample of control participants; neither patient significantly differs from the controls (Crawford-Howell single-case test *p*s>0.499). Similarly, in the matrix reasoning task, both patients solved the vast majority of the problems (25/30 and 26/30), which puts them +2.3 and +1.8 standard deviations above the mean based on the age-matched normative data available for this task (WASI-II; (109)) (these raw scores correspond to *T*-scores of 73 and 68; normative *T* = 50, SD = 10).

### Logical reasoning elicits strong responses outside the language network

Given that logical reasoning demands in our paradigms did not engage the language network, we wanted to ensure that they elicit a response somewhere in the brain. We first examined neural responses in one plausible candidate system—the Multiple Demand (MD) network (51, 132, 133). This network comprises a set of bilateral frontal and parietal brain areas that show strong activity during diverse cognitively demanding tasks, such as standard executive function tasks (132, 134–137). Damage to the MD network is associated with decreases in fluid intelligence (138–140). In addition, and most relevantly here, certain forms of reasoning recruit the MD network, including mathematical reasoning (40, 41, 141) and the kind of reasoning necessary to understand computer code (42, 43, 142). Replicating much prior work (e.g., (141, 143)), the areas of the MD network showed a robust response to a spatial working memory task, including a stronger response during the more demanding condition estimated in independent data (*p*<0.001 at the network level; **SI Figure 2** and **SI Table 2**). The inductive reasoning task elicited a robust response across the MD network, with a stronger response during the rule induction condition compared to the control, rule application condition (*p*<0.001 at the network level; **SI Figure 2**, **SI Table 2**). So did the matrix reasoning task (*p*<0.001 at the network level; **SI Figure 2, SI Table 2**). However, the deductive syllogistic reasoning task did not engage the MD network (*p*>.48 at the network level). This finding aligns with the results in Coetzee & Monti (2018), who reported a dissociation between brain areas sensitive to deductive reasoning versus to general task difficulty, the latter being a key signature of the MD network (see also (57, 144, 145)). A whole-brain search revealed several frontal and parietal areas that showed a stronger response to the Modus Tollens condition compared to the Modus Ponens condition estimated in independent data (*p*<0.001 across the set of areas; **SI Figure 2**; see (146) for a detailed investigation of these brain areas).

## Discussion

To shed light on the long-standing debate on the role of linguistic representations in formal logical reasoning, we examined the responses of the language brain areas—robustly sensitive to linguistic syntactic structure (29, 32, 36, 37)—to inductive and deductive reasoning. In a complementary approach, we tested logical reasoning abilities in individuals with severe damage to the language areas. The results converged on a clear answer: linguistic representations are not engaged nor necessary for logical reasoning, at least in the adult brain. Below, we contextualize these findings with respect to the broader literature.

### Accumulating evidence for the separability of language from other cognitive functions

Our results showing that the left-lateralized language network is not engaged in logical reasoning add to a growing body of evidence that the language network supports specifically linguistic computations. Relative to our perceptual and motor systems, our memory and attention mechanisms, and our social reasoning capacities, language is a relatively recent cultural invention (147). As a result, it is reasonable to hypothesize that when language emerged, it co-opted various components of pre-existing brain machinery. However, although language undeniably shares similarities with diverse non-linguistic functions, hypotheses about neural overlap have not found empirical support. For example, despite the fact that language serves a social-communicative function (148–150, 27) and is similar, in some ways, to non-verbal socially-relevant signals, such as facial expressions, non-verbal vocalizations, and gestures, evidence suggests that such non-verbal signals are processed in brain areas distinct from the language network (e.g., (151–153)). Despite the fact that language shares hierarchical structure with music and other domains (154, 155), music does not engage the language areas (39, 156, 157). And despite the fact that language processing requires executive resources, such as working memory and cognitive control (158–162), executive tasks do not engage the language network (39, 48, 49), and demanding linguistic computations—for example, forming non-local syntactic dependencies—are instead carried out locally within the language network ((34, 36, 163–165) see (166) for a review).

Of greatest relevance to the current investigation, the syntactic structure of language bears similarity to structure that characterizes abstract reasoning, including formal—mathematical and logical—thinking (11, 167, 168), as well as domain-specific reasoning, such as intuitive physical reasoning (169–171) and social reasoning (172). However, as with other non-linguistic functions, the studies that had examined the relationship between language processing and these types of reasoning found no overlap: mathematical reasoning (39, 41, 173) (for evidence from aphasia, see (22)), computer code comprehension (42, 43), physical reasoning (174), and social reasoning (44–46, 175) (for evidence from aphasia, see (23, 25, 176)). Our study adds to this body of work, showing that formal logical reasoning—including both induction and deduction—does not recruit nor require linguistic representations.

### Challenges to the view that language does not support thinking?

Some have challenged the separability of language and thought based on correlations between the degree of linguistic impairment and impairment on certain cognitive tasks (e.g., (52–55)). However, these effects can be explained by the proximity between the language areas and other cognitive areas (125, 177, 178)—and, in some cases, co-lateralization of those functions to the left hemisphere (e.g., language and arithmetic: (179))—combined with the fact that stroke damage does not respect functional boundaries. For this reason, dissociations are generally considered more informative than associations (180).

Another potential challenge to the language-thought separability comes from dual-task studies that show that a verbal task can interfere with one’s ability to perform a non-verbal cognitive task (e.g., (181–183)). However, the empirical landscape is complex (for a review, see (56)), and many studies do not meet the evidential standard necessary to argue for the role of verbal representations in non-linguistic cognition. In particular, to argue that a verbal task that interferes with some critical task shares resources with that task, it is important to include a control interference task, matched for difficulty to the verbal task (evaluated in a single-task design), and to obtain an interaction such that the verbal task affects performance on the critical task to a reliably greater extent; the non-verbal task should further be shown to reliably affect performance on some other task (see (184) for a discussion of these methodological issues). In addition to these design and statistical considerations, many verbal interference paradigms (e.g., syllable or word/nonword repetition tasks) engage lower-level speech articulation and perception areas (e.g., (185, 186)), not the language areas. Because most proposals about the role of language in thinking concern lexical and/or syntactic representations (which are handled by the higher-level language areas; (38)), it is difficult to know what verbal interference effects—even in studies that are properly designed—would mean for these proposals.

Thus, to our knowledge, no prior study—based on neuropsychological patient data or using verbal interference paradigms—provides unambiguous evidence that humans rely on lexical or syntactic representations to think, but we remain open to seeing such studies in the future, including studies that may show transient recruitment of linguistic representations, which may not be easily detectable with fMRI but may show up in studies that use approaches such as intracranial recordings.

### Neural dissociation does not entail encapsulation

The evidence that language and thinking are dissociable in the human mind and brain is perfectly compatible with the fact that the language network has to constantly interact with our knowledge and reasoning systems (see (38, 90, 187) for discussion). First, language is only useful insofar as it allows us to share ideas with one another: both language production and comprehension therefore necessarily require our knowledge and reasoning systems in addition to our core linguistic mechanisms. Moreover, language is a powerful information compression tool, allowing for efficient representation of concepts that are useful to a given culture (188–194). In this way, language may play a facilitatory role during some cognitive tasks, but not because we use words and linguistic syntax to represent and manipulate ideas, but because using words or non-linguistic symbols (e.g., Arabic digits) or schematics (e.g., Venn diagrams) allows offloading partial or intermediate products of internal computation. Whether language is special relative to other compression tools is not known and would require careful comparisons. Finally, language can certainly facilitate learning of certain ideas, including abstract logical or relational structures (e.g., (195–199, 67, 200)). However, this role of language in concept acquisition again does not imply that we use linguistic representations to think any more than learning concepts from visual inputs implies that we use our visual system to think.

### Why do language processing and logical reasoning dissociate?

The language-logic dissociation presumably stems from the distinct demands associated with linguistic communication vs. formal reasoning. One key difference may have to do with the kinds of meanings these systems express: natural languages tend to express meanings related to the external and internal world, but mathematics, logic, and computer code express mostly abstract, relational meanings that do not bear a direct or necessary connection to the world (see (201) for discussion). This difference may provide a strong signal during the early processing of the input that a particular problem should be routed to a particular system. Of course, if a problem is cast into a sentence form (as in our syllogistic reasoning task here), it necessarily has to go through the language system to be parsed, before it can be solved by a (putatively downstream) reasoning system.

Another reason for the dissociation may be that the linguistic format is not well-suited for formal reasoning: in spite of superficial similarities in the structured nature of linguistic and logical expressions, linguistic representations are noisy and ambiguous in ways that make them unreliable for supporting formal inference. Natural language is riddled with referential vagueness, scope ambiguities, and underspecification, requiring pragmatic enrichment, all of which can obscure the logical structure of a sentence. Indeed, recent evidence from AI research suggests that language-only neural network models can struggle with formal reasoning tasks (202–204, 90, 92), but converting linguistic inputs into logical formalisms (e.g. statements in first-order logic), which can then be fed into an external theorem solver, leads to large performance gains (e.g., (205–215)).

### Alternatives to linguistic representations

If we don’t think in language, what kind of representations support logical reasoning? Competing accounts postulate distinct formats of thought-mediating representations, which could translate into distinct predictions about neural implementation. For example, according to the *mental model* framework, reasoners construct analog simulations of the problem, exploiting visuo-spatial representations and working memory resources to model the possible consequences of premises (216, 217). These accounts therefore predict the engagement of brain areas that support visual/spatial imagery and working memory (218, 219). Alternatively, *script– or schema-based* approaches propose that reasoning draws on cached relational templates (structured representations of events or situations abstracted from prior experience) (220–223). These accounts suggest that reasoning is scaffolded by context– and domain-specific knowledge structures, which may draw on multiple semantic networks (e.g., (224, 225)). And the *mental logic* proposals construe abstract reasoning as the manipulation of amodal, abstract symbols according to syntactic inference rules (226–228). Perhaps the most promising candidate from the latter class of proposals is the probabilistic language of thought (PLOT) hypothesis (229–235). According to this hypothesis, mental algorithms are construed as symbolic programs over concepts, which encode probabilistic knowledge, including both domain-specific knowledge and abstract relational knowledge. In this way, the PLOT provides a flexible medium for any type of reasoning, and captures the graded nature of mental representations. Future brain imaging studies may help determine whether logical reasoning depends on visuo-spatial, fully abstract, or domain-specific (at least during early stages of development) representations.

### Modularity of reasoning

Aside from the language-logic dissociation, this study highlights the fact that different kinds of reasoning dissociate from one another. Some forms of reasoning have been shown to be domain-specific, including social reasoning (236–239) and intuitive physical reasoning (240–243). Both of these systems are distinct from the Multiple Demand (MD) network, implicated in diverse goal-directed behaviors and some forms of abstract reasoning (51, 137), and from the Default network, implicated in episodic self-projection and constructing situation models (244–247), perhaps using spatial representations (248). We here show that inductive and matrix reasoning, similar to arithmetic reasoning, strongly engage the MD network, but syllogistic deductive reasoning does not, and instead recruits a distinct set of brain areas, in line with some past studies ((57, 101, 144, 145); see (146) for a systematic investigation of these areas). Why do these different forms of reasoning dissociate? Is it because of the distinct content/structure of the problems? Or, perhaps, differences in the mental operations that those problems require? Given the strong integration between memory and computation in the brain (see e.g., (249) and (250) for discussions), both of these factors may contribute to the observed dissociations.

### Limitations and open questions

The current study is limited in at least two ways. (#1) The aphasia component includes only two participants. This small number was dictated by the requirement of a *profound* linguistic impairment, which is rare (251). Evidence of intact logical abilities in patients with less severe aphasia would be less informative because it would always be possible that the patients are relying on the remaining portions of the language network to reason. Evidence of intact cognition in severely linguistically impaired individuals is precious and important, even if it comes from a handful of participants. After all, the field of cognitive neuroscience has gained some of its greatest insights from case studies (252–255). (#2) These findings should be extended to a broader array of logical reasoning tasks, including contextualized paradigms (256–259), and non-verbal versions of purely deductive tasks.

Many questions remain open. For example: Is the language-logic dissociation already present during early childhood, or does it emerge over development? Is linguistic exposure necessary to learn to reason logically? And is the dissociation between language and logic, or among different types of reasoning, an inevitability given the computational demands of these different cognitive tasks, or merely an accident of biological evolution—a question that can now be tested in neural network models (260–264).

### Conclusions

Our results provide evidence against the hypothesis that natural language serves as the medium of abstract logical reasoning and suggest that such reasoning is underpinned by a distinct representational format.

## MATERIALS AND METHODS

### Study 1 (fMRI) Participants

Twentynine adults (15 females; mean age=27.3 years, SD=12.9 years) from MIT and the surrounding Boston, MA community participated for payment. All were native English speakers, had normal hearing and vision, no history of language impairment, and canonical left-lateralized language networks (as determined by the language localizer). All participants provided written informed consent in line with the requirements of MIT’s Committee on the Use of Humans as Experimental Subjects (Protocol #2010000243). All participants completed the language localizer task and the Multiple Demand (MD) localizer task; different subsets of the 29 participants completed the logic tasks, with at least 17 participants per task (17 completed the induction and the matrix reasoning tasks; 23 completed the syllogistic reasoning task).

### Study 2 (behavior) Participants

Two profoundly aphasic male participants participated (S.A.: 78 years; G.S.: 50 years). Both had large lesions that had significantly damaged the left inferior frontal and left temporal brain areas, and were classified as severely agrammatic (**SI Table 3**). Both were native English speakers, did not present with any visual impairments, and were pre-morbidly right-handed. Forty age-matched neurotypical participants were included as controls (see <u>Supp. Methods</u> for details). Ethics approval was granted by the UCL Language and Cognition Departmental Ethics Committee (LC/2023/05). All participants provided informed consent prior to taking part in the study.

### Inductive Reasoning Task (fMRI, behavior)

Participants were shown an input list of 1-5 single-digit numbers and an output list of 0-5 single-digit numbers and asked to guess the rule that transformed the input list into the output list. They were then shown another input list and asked to type the output list. They were told whether or not their guess about the underlying rule is correct and are shown another input list. In the **fMRI version**, each trial (corresponding to a rule) consisted of 8 problems. After the last problem, participants were told—via a schematic (**Figure 1**, left)—the correct rule and asked to apply this rule to two more input lists. We defined the *induction* period as the 8 problems before the correct rule was revealed, and the *application* period as the two problems after the correct rule was revealed. The rule induction > rule application contrast targets neural processes involved in hypothesis generation and rule discovery beyond those required for rule implementation and response entry. Each participant completed 40 rules. For this and other tasks, all the materials are available at OSF (https://osf.io/jm9rd); additional procedural and timing details are included in <u>Supp. Methods</u>.

For the **behavioral version** of the task, the problems were presented on paper, and participants provided written responses. The same 40 rules were used, except that a few problems were added in order to have 14 problems (input-output pairs) per rule, and a few problems were replaced based on feedback from the fMRI participants who found some input-output pairs confusing. The stimuli were divided into four sets, with ten rules per set. GS completed all 40 rules across several sessions (∼6 rules per session); SA only completed sets 1-3 also across sessions (∼4 rules per session); the variability in the number of rules per session was due to age-related health conditions and because the experiment had to be completed amongst other activities at routine check-in sessions. Control participants each completed one set of 10 rules within a single testing session, with a brief break half-way through, and 10 participants completed each set. For both the patients and the controls, testing for a given rule was stopped after the participant answered four problems in a row correctly. Time permitting, the participants with aphasia were sometimes offered to schematically illustrate the rule they guessed (see **Figure 2** for examples).

### Deductive (Syllogistic) Reasoning Task (fMRI)

Participants were presented with classic syllogisms and asked to judge the validity of the third sentence (the conclusion) given the first two sentences (the premises). Each trial corresponded to a syllogism, and trials varied in difficulty between harder syllogisms (Modus Tollens) and easier syllogisms (Modus Ponens), and in whether the problems used real words or nonwords. The hard syllogisms > easy syllogisms contrast targets cognitive processes related to the complexity of logical deductive reasoning. Each participant completed between 16 and 32 trials of each condition.

### Matrix reasoning task (fMRI, behavior)

Participants were presented with sets of images of geometrical shapes and asked to find an outlier (fMRI version) or a missing image (behavioral version). In **fMRI**, participants were presented with four images and asked to decide which image does not fit with the others. The experiment used a blocked design; trials in the hard blocks used stimuli from Cattell (107), and trials in the easy blocks used simpler problems created by Woolgar et al. (141), who adapted this task for fMRI (e.g., three images of the same simple shape and an image of a different shape); this easier condition can be solved by visual pattern recognition. The hard patterns > easy patterns contrast targets cognitive processes related to relational reasoning, hypothesis generation, and deductive inference.

For the **behavioral** version, we used the Matrix Reasoning subtest of the Wechsler Abbreviated Scale of Intelligence, Second Edition (WASI-II). Participants were presented with an array of images with one image missing and asked to choose the missing image from five options to complete the pattern. The test was administered on paper, following the standard WASI-II administration procedures. Testing was stopped after the participant answered three problems incorrectly.

### Critical Analyses – fMRI

(For the details of fMRI data acquisition, preprocessing, and modeling, see <u>Supp. Methods</u>.) To test whether the language areas respond during logical reasoning tasks, we used an extensively validated localizer (122) to identify regions of interest functionally in individual participants. Participants were asked to attentively read sentences and sequences of non-words (for details, see <u>Supp. Methods</u>). The responses in the language fROIs (and control fROIs, as explained below) were statistically evaluated using linear mixed-effects models. For each contrast, we fit separate models at both the network level and for individual fROIs.

The **network-level model**: *BOLD ∼Condition + (1 | Participant) + (1 | fROI)*.

The **individual fROI-level model**: *BOLD ∼Condition + (1 | Participant)*.

To ensure that our logical reasoning tasks elicit a strong response somewhere in the brain, we performed two analyses. *First*, we used a spatial working memory task (keeping track of 8 vs. 4 squares in a 3×4 grid) as a localizer for the Multiple Demand network (e.g., (143)) (for details, see <u>Supp. Methods</u>). And *second*, because the deductive syllogistic reasoning task did not elicit a response in the MD network, we additionally performed a whole-brain search for areas that respond to the hard syllogisms > easy syllogisms contrast (see <u>Supp. Methods</u>). (For information on the behavioral performance of participants in the critical fMRI tasks, see **SI Figure 4**)

### Critical Analyses – Behavior

For the inductive reasoning task, we used the Crawford and Howell (265) test to compare each patient’s performance to the controls. The Matrix Reasoning WASI-II data were scored according to the standard guidelines provided in the official manual.

## Acknowledgements

We would like to acknowledge the Athinoula A. Martinos Imaging Center at the McGovern Institute for Brain Research at MIT, including the technical team—Steve Shannon and Atsushi Takahashi. We would also like to thank Anya Ivanova, Colton Casto, Ben Lipkin, Chiebuka Ohams, and Ted Gibson for helpful comments on the manuscript. HK was supported by a Friends of McGovern graduate fellowship and a graduate fellowship from the K. Lisa Yang Integrative Computational Neuroscience (ICoN) Center. JBT was supported by research funds from the Air Force Office of Scientific Research, the Office of Naval Research Science of AI program, and a Schmidt Sciences AI2050 Fellowship. JSR and STP were supported by the Division of Research on Learning in Formal and Informal Settings (EDU/DRL) grant 2201843. RV was supported by a Leverhume Research Fellowship (RF-2023-690\10). EF and this research were partially supported by research funds from the McGovern Institute for Brain Research, the Department of Brain and Cognitive Sciences, MIT’s Siegel Family Quest for Intelligence, a grant from the Simons Founda-tion to the Simons Center for the Social Brain at MIT, and a grant from the DARPA AIQ program through the DARPA CMO contract number HR00112520025.

---

**Supplementary Figure 1.**
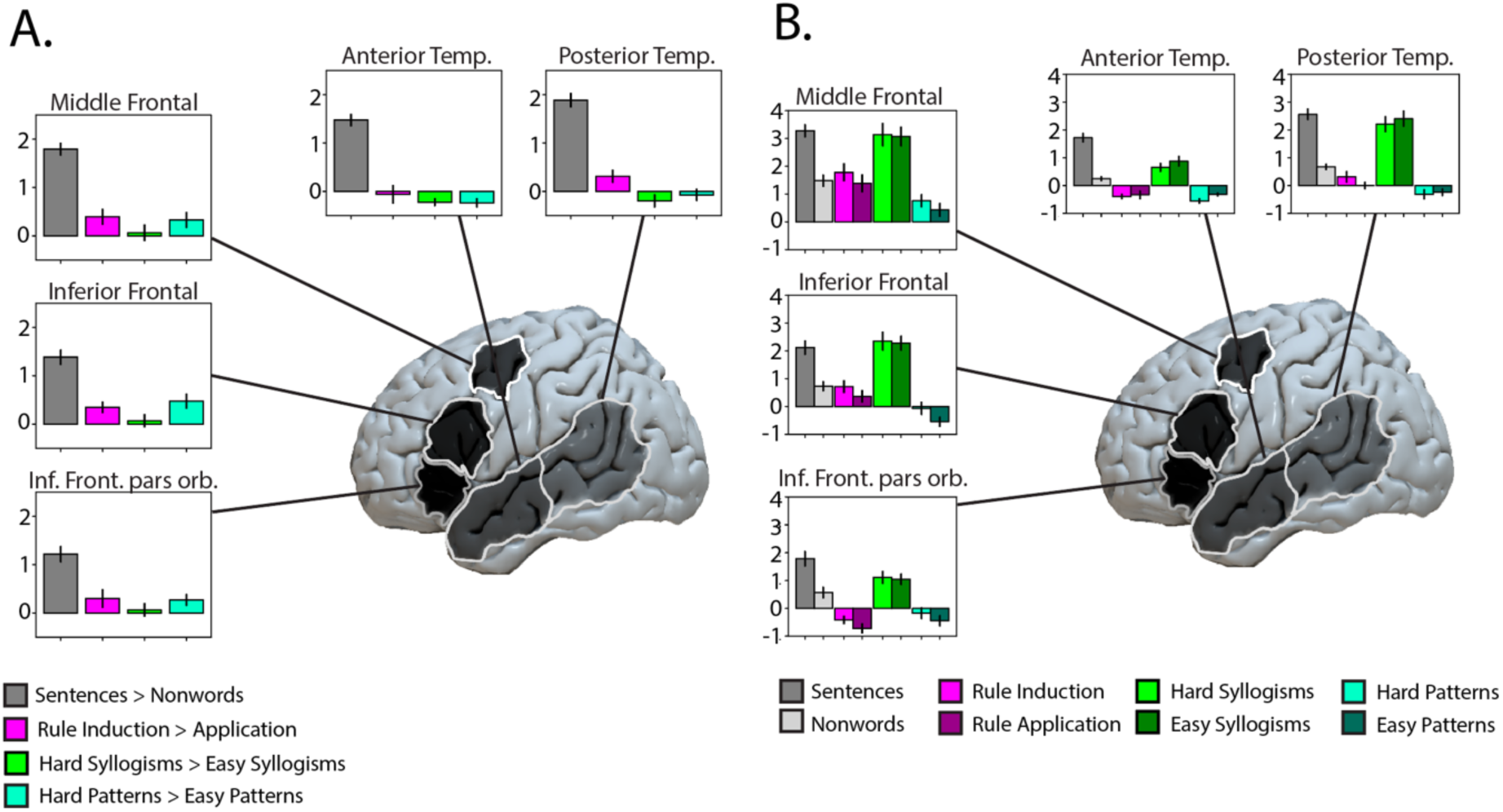
The language network does not support logical reasoning: The results broken down by fROI. **A.** Responses in the five language fROIs (individually defined and constituting 10% of the broad masks shown in grey on the brain; see <u>Methods</u>), in percent BOLD signal change, to the language (grey), inductive reasoning (magenta), deductive reasoning (green), and matrix reasoning (teal) contrasts. The style matches the style of main Figure 2B. **B.** Responses in the five language fROIs to the individual conditions relative to the low-level fixation baseline (cf. panel A, which shows the differences between the two conditions for each experiment).

**Supplementary Figure 2.**
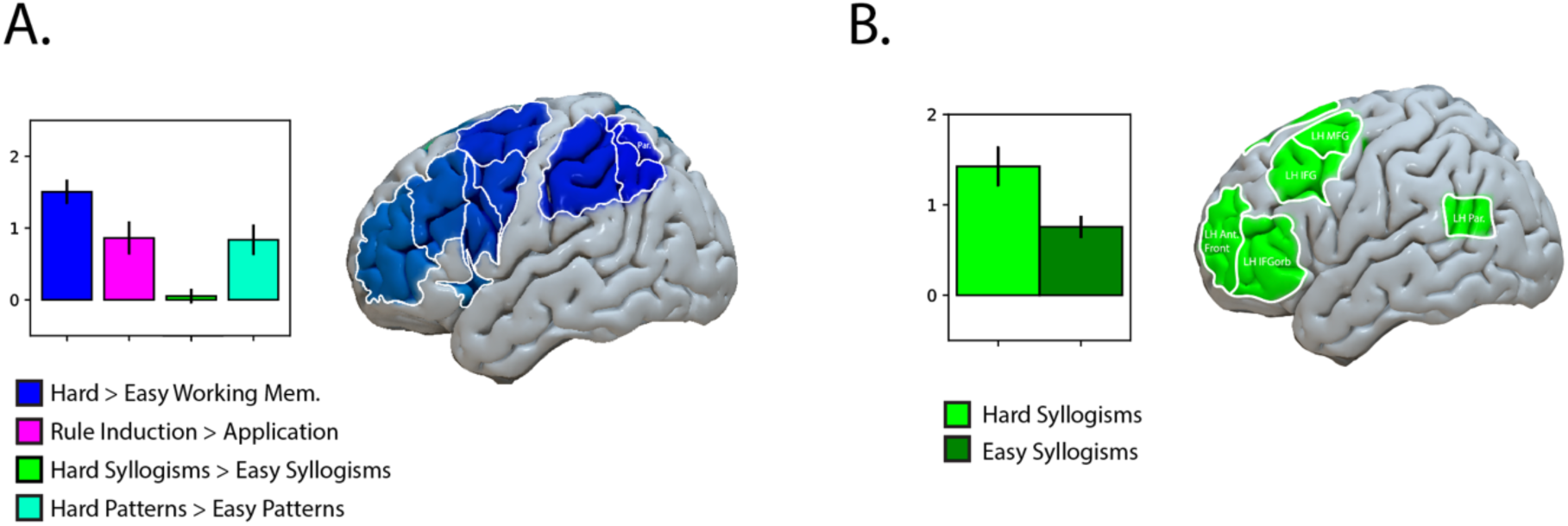
The brain networks that support logical reasoning. **A.** Responses in the Multiple Demand (MD) network, in percent BOLD signal change, to the MD localizer contrast (a spatial working memory task; see <u>Methods</u>; blue) and the three logical reasoning contrasts (magenta, green, and teal). The MD fROIs are defined within individuals and constitute 10% of the broad masks shown in blue on the brain (see <u>Methods</u>). The MD network shows a reliable response to the inductive reasoning and the matrix reasoning contrasts, but not to the deductive (syllogistic) reasoning contrast. **B.** The brain areas identified with the Hard Syllogisms > Easy Syllogisms contrast. The fROIs are defined within individuals (using a portion of the data) and constitute 10% of the broad masks shown in green on the brain. The responses are estimated using left-out data and reveal a robust and replicable effect.

**Supplementary Figure 3.**
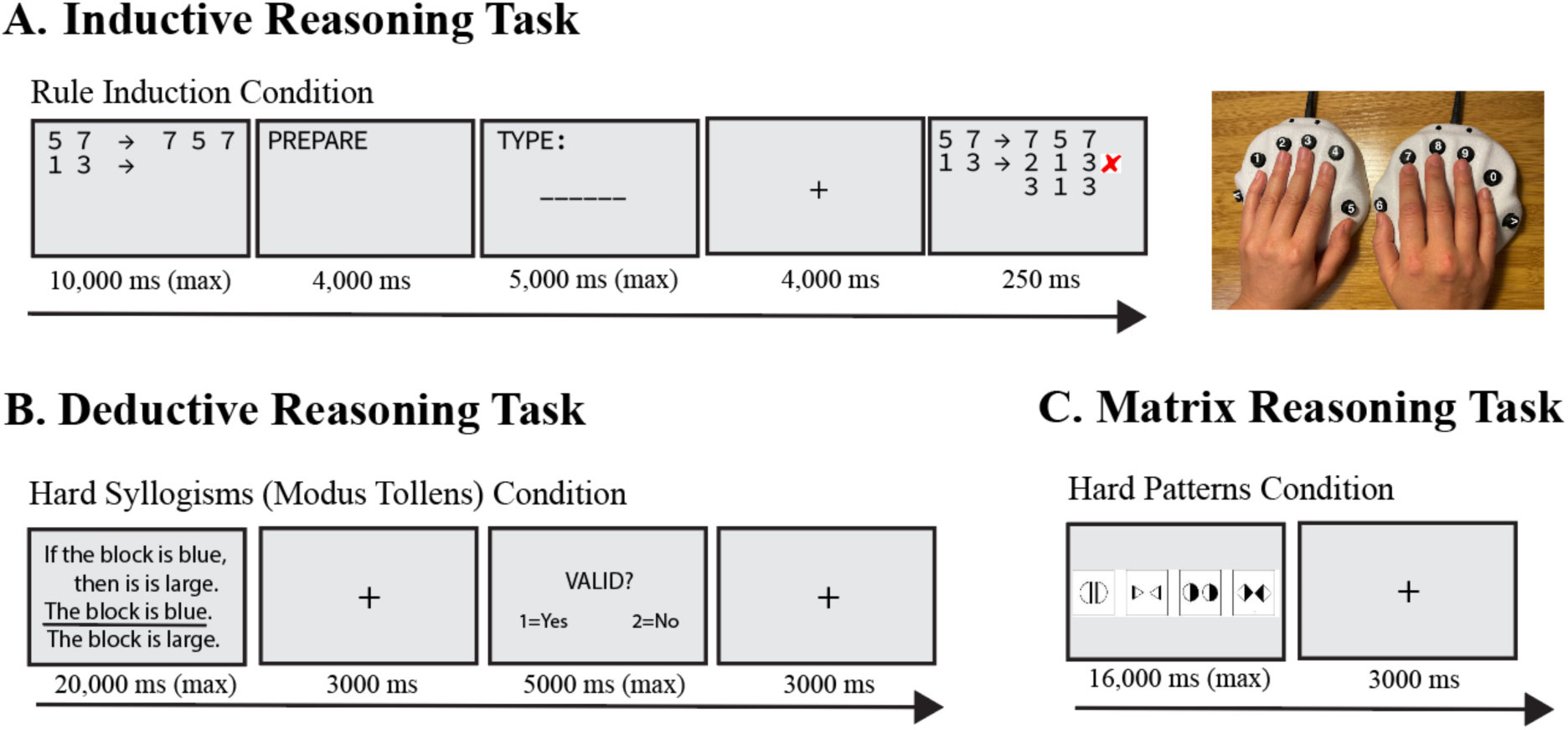
Timing details for the three reasoning tasks used in the fMRI study. **A.** A sample trial for the Inductive Reasoning Task, in which participants iteratively generated outputs from given input lists and received trial-by-trial feedback to discover an underlying transformation rule. Each rule block consisted of 8 problems followed by 2 rule application trials after the rule was revealed. Blocks were self-paced, and each scanning run included 2 rule blocks. **B.** A sample trial from the Hard Syllogisms condition of the Deductive Reasoning Task, in which participants judged the validity of the conclusion given the premises. Trials were self-paced, and each run included 16 trials. **C.** A sample trial from the Hard Patterns condition of the Matrix Reasoning Task, in which participants had to figure out the rule that applies to the images and find the outlier image. Trials were self-paced and grouped into 16-second blocks, and each run included 16 blocks. (See OSF (https://osf.io/jm9rd) for all task materials.)

**Supplementary Figure 4.**
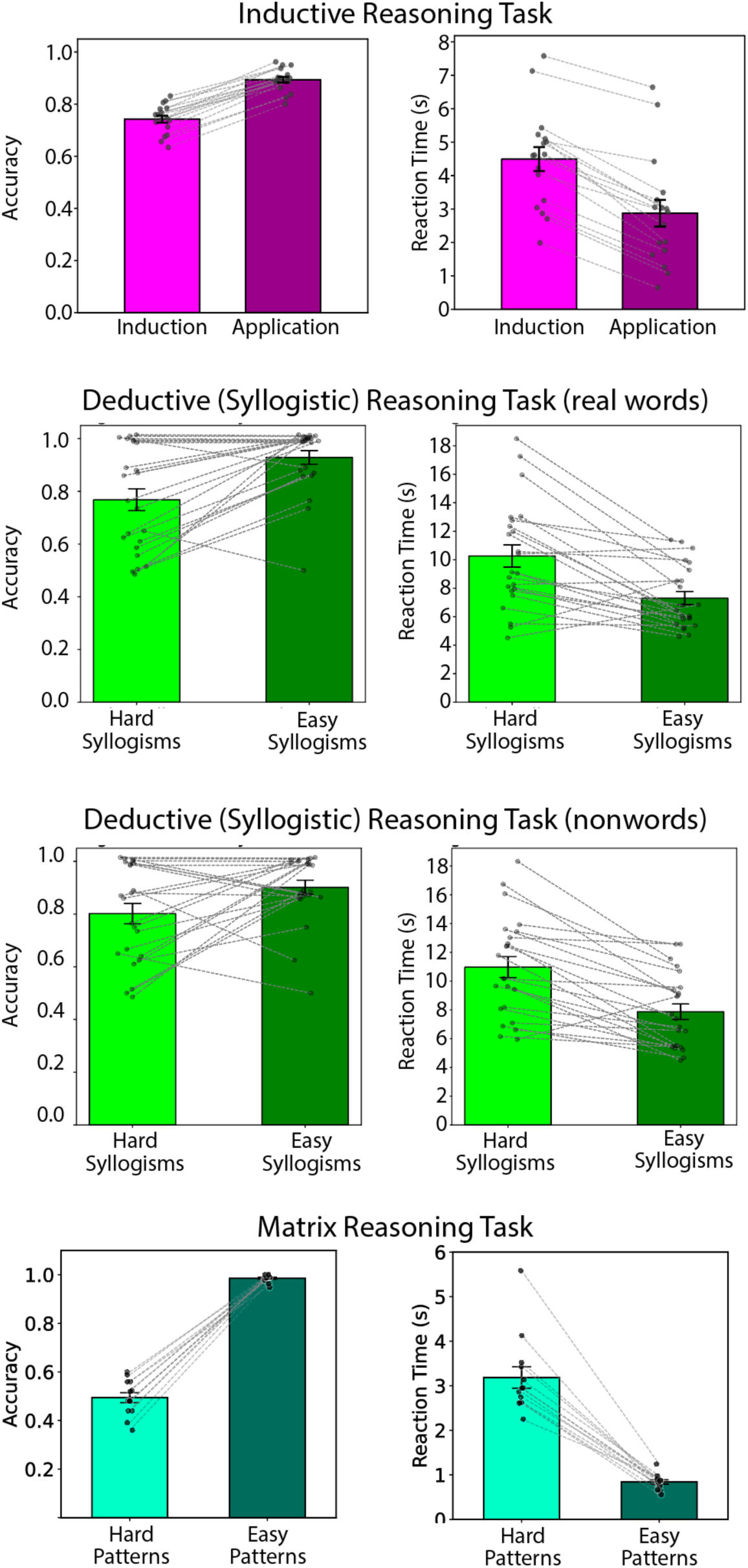
Behavioral performance for the reasoning tasks. Accuracies and reaction times (RTs) for the inductive, deductive, and matrix reasoning tasks. For the deductive (syllogistic) reasoning task, we break down the trials by whether they used real words or nonwords. For all graphs, the error bars correspond to the standard error of the mean by participant, the dots correspond to individual participants’ responses, and the thin dashed lines connect the same participants across conditions. The overall response rate was 100% for the induction task, 99.63% for the deductive syllogistic reasoning task, and 100% for the matrix reasoning task. The reaction times (RTs) mirrored the accuracy data, with longer RTs for the harder conditions.

**Supplementary Figure 5.**
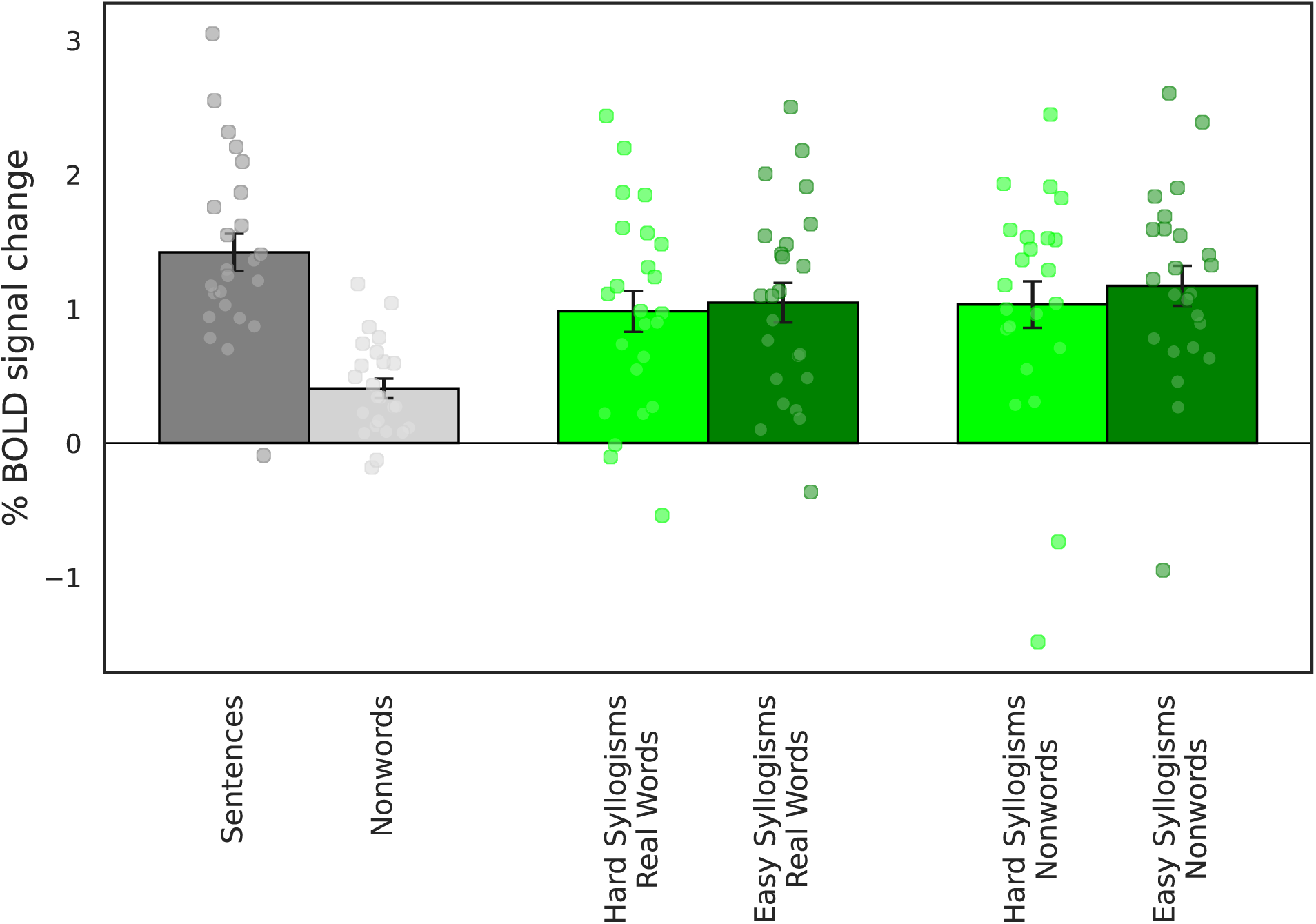
The language network’s responses to the deductive (syllogistic) task broken down by real word vs. nonword syllogisms. Responses in the language network, in percent BOLD signal change, to the conditions of the language localizer (estimated in left-out data) and to the conditions of the deductive syllogistic reasoning task broken down by type of stimulus (words vs. nonwords). The error bars correspond to the standard error of the mean by participant, and the dots correspond to individual participants’ responses.

**Supplementary Table 1.** The responses of the language network to the language and logical reasoning contrasts broken down by fROI. The results from the linear mixed-effects models (see <u>Methods</u>) at the level of individual language fROIs. We report uncorrected p-values, and we mark the values that survive the Bonferroni correction for the number of fROIs (n=5) in **bold**.

| Task | fROI | Estimate | p-value |
| --- | --- | --- | --- |
| Language | LH Inferior Frontal Gyrus (orbital) | **1.218** | **<0.0001** |
| Language | LH Inferior Frontal Gyrus | **1.386** | **<0.0001** |
| Language | LH Middle Frontal Gyrus | **1.793** | **<0.0001** |
| Language | LH Anterior Temporal | **1.476** | **<0.0001** |
| Language | LH Posterior Temporal | **1.887** | **<0.0001** |
| Induction | LH Inferior Frontal Gyrus (orbital) | 0.302 | 0.1360 |
| Induction | LH Inferior Frontal Gyrus | 0.350 | 0.0092 |
| Induction | LH Middle Frontal Gyrus | 0.396 | 0.0301 |
| Induction | LH Anterior Temporal | -0.060 | 0.7500 |
| Induction | LH Posterior Temporal | 0.312 | 0.0356 |
| Deduction | LH Inferior Frontal Gyrus (orbital) | 0.0647 | 0.6466 |
| Deduction | LH Inferior Frontal Gyrus | 0.0702 | 0.6059 |
| Deduction | LH Middle Frontal Gyrus | 0.0645 | 0.7105 |
| Deduction | LH Anterior Temporal | -0.225 | 0.0117 |
| Deduction | LH Posterior Temporal | -0.197 | 0.1717 |
| Matrix | LH Inferior Frontal Gyrus (orbital) | 0.272 | 0.0448 |
| Matrix | LH Inferior Frontal Gyrus | 0.477 | 0.0077 |
| Matrix | LH Middle Frontal Gyrus | 0.328 | 0.0655 |
| Matrix | LH Anterior Temporal | -0.238 | 0.0265 |
| Matrix | LH Posterior Temporal | -0.073 | 0.5734 |

**Supplementary Table 2.** The responses of the Multiple Demand network to the spatial working memory and logical reasoning contrasts, and of the deduction-sensitive areas to the deductive reasoning contrast. The results from the linear mixed-effects models (see <u>Methods</u>) at the network level.

| Multiple Demand Network |  |  | Deduction-sensitive fROIs |  |  |
| --- | --- | --- | --- | --- | --- |
| Predictors | Betas | p-value | Predictors | Betas | p-value |
| *Spatial WM Task* |  |  | *Deductive Reasoning Task* |  |  |
| Intercept (easy) | 1.83 | <0.0001 | Intercept (MP) | 0.76 | 0.0016 |
| Critical (hard) | 1.50 | <0.0001 | Critical (MT) | 0.67 | <0.0001 |
| *Inductive Reasoning Task* |  |  |  |  |  |
| Intercept (application) | 1.81 | <0.0001 |  |  |  |
| Critical (induction) | 0.86 | <0.0001 |  |  |  |
| *Deductive Reasoning Task* |  |  |  |  |  |
| Intercept (Modus Ponens) | 2.26 | <0.0001 |  |  |  |
| Critical (Modus Tollens) | 0.05 | 0.4822 |  |  |  |
| *Matrix Reasoning Task* |  |  |  |  |  |
| Intercept (easy patterns) | 1.67 | <0.0001 |  |  |  |
| Critical (hard patterns) | 0.84 | <0.0001 |  |  |  |

**Supplementary Table 3.** Extended demographic and language assessment information for participants with aphasia. G.S. was premorbidly in cybersecurity, and S.A. was premorbidly a police sergeant. GS suffered a left-hemisphere stroke, also resulting in a vascular lesion in the left middle cerebral artery and damage to the perisylvian areas. S.A. suffered a subdural empyema, followed by meningitis, leading to a secondary vascular lesion in the left middle cerebral artery and damage to the perisylvian areas. This table further shows cognitive assessments include the WAIS digit span (1), Pyramid and Palm Trees test (3-picture version), identifying semantic relations between pictures (2), and WASI matrices (3). Grammatical assessments include comprehension of spoken and written role-reversible sentences (materials from (4)) and grammaticality judgments (materials from (5)), with repetition penalties for spoken/auditory versions. Lexical assessments include the spoken and written picture matching and synonym judgement tasks from the ADA comprehension battery (6) and the picture naming task from the PALPA test battery (7). * indicates performance above chance under a binomial test at p = 0.05. ** a cut-off was applied for SA’s PALPA naming test: with 1/20 on the high frequency set, the mid– and low-frequency sets were automatically scored 0/20 without further testing.

|  | GS | SA |
| --- | --- | --- |
| **Age** | 50 | 78 |
| **Time post-onset** | 2.5 years | 33 years |
| **Etiology** | Left hemisphere stroke | Left subdural empyema and meningitis |
| **Previous occupation and educational background** | Graduate degree; worked in cybersecurity | Left school at 15; police sergeant |
| **Cognitive assessments**<br>- Forward digit span (Wechsler Adult Intelligence Scale)<br>- Pyramid & Palm Trees (3-picture version) | - 3 items<br>- 50/52 | - 2 items<br>- 49/52 |
| **Grammatical processing**<br>Comprehension spoken reversible sentences:<br>- Total (chance = 40/80)<br>- Active (20/40)<br>- Passive (20/40)<br>Comprehension written reversible sentences:<br>- Total (chance = 40/80)<br>- Active (20/40)<br>- Passive (20/40)<br>Auditory grammaticality judgment:<br>- Total (chance = 32/64)<br>Written grammaticality judgment:<br>- Total (chance = 32/64) | - 22/80<br>- 20/40<br>- 2/40<br>- 38/80<br>- 32/40*<br>- 6/40<br>- 36/64<br>- 44/64* | - 30/80<br>- 16/40<br>- 14/40<br>- 46/80<br>- 34/40*<br>- 12/40<br>- 48/64*<br>- 29/64 |
| **Lexical processing**<br>ADA spoken-word picture matching (total):<br>ADA written-word picture matching (total):<br>PALPA Picture Name (spoken or written):<br>- High-frequency set<br>- Mid-frequency set<br>- Low-frequency set<br>ADA Synonym Judge (combined written and auditory presentation):<br>- Total<br>- High imageability errors<br>- Low imageability errors<br>- High frequency errors<br>- Low frequency errors | - 60/66<br>- 63/66<br>- 19/20<br>- 15/20<br>- 13/20<br>- 137/160<br>- 2<br>- 10<br>- 4<br>- 7 | - 51/66<br>- 53/66<br>- 1/20<br>- 0/20**<br>- 0/20**<br>- 135/160<br>- 4<br>- 12<br>- 6<br>- 3 |

## Supplementary Methods

### 1. Additional participant details

#### Study 1

26 participants were right-handed, 1 was ambidextrous, and 2 were left-handed. Critically, all participants had left-lateralized language networks, as determined by the language localizer task.

#### Study 2

The mean age of the 40 control participants (20 female) was 56 years (SD = 10.7, range: 40 to 86 years). The mean number of years of education was 17.2 (SD = 2.3), perhaps driven up by the fact that our study was advertised as consisting of logic puzzles. All participants were native English speakers, had normal, or corrected-to-normal vision, and no history of language impairment, neurological diseases, or reading impairments, and performed within the normal range on the Mini Mental State Examination (MMSE; (8)). Both the patients with aphasia and the control participants completed the experiments individually, in a quiet room, with an experimenter present throughout the testing session.

### 2. Additional details of the critical paradigms

#### Inductive Reasoning Task

The rules were distributed across 20 scanning runs, with 2 rules per run and each run lasting between 200s and 420s (durations are variable given the self-paced nature of the task; see **SI Figure 3A** for the timing details). Participants entered responses via a scanner-safe fiber-optic button response device (Nata Technologies LxPad system), which contains 12 buttons allowing input of digits 0–9, backspace, and enter (**Figure 1**, left panel).

For the behavioral version, participants with aphasia were shown an example rule prior to the experiment, which acted as a training item. The order of the rules within a set was kept constant across participants, but the order of the problems within a rule was randomized.

#### Deductive Reasoning Task

The trials were distributed across two scanning runs, with 8 trials per condition per run and each run lasting between 551 s and 1,034 s (durations are variable given the self-paced nature of the task; see **SI Figure 3B** for the timing details). Participants responded by pressing one of two buttons on the button box. The experiment additionally included two conditions of no interest (challenging memory conditions).

#### Matrix Reasoning Task

In fMRI, each participant completed a single run of the task lasting 320 s and consisting of 8 easy blocks and 8 hard blocks. The blocks were of fixed length (16 s), which means that only a few hard trials could be solved during this period (between one and six), and more easy trials could be solved (between 4 and 15; see **SI Figure 3C** for the timing details); furthermore, because only 24 hard items were available, some participants did not have enough items for all 8 blocks, but every participant completed at least 5 hard blocks. Participants responded by pressing one of four buttons on the button box.

For the behavioral version, the items (total number = 30) were presented in ascending order of difficulty. Prior to the main test, participants were shown a short set of practice items to familiarize them with the task. These practice items illustrated the nature of the patterns, and participants were provided with explanations (including gestural cues, where necessary) to aid comprehension. Participants indicated their choice by pointing. For scoring and analyzing the matrix reasoning task, each item was evaluated for accuracy, and raw scores (total correct out of 30) were converted into *T*-scores using age-normed tables (μ = 50, σ = 10). These *T*-scores were the basis for statistical comparison with the normative control sample from the testing manual, consistent with standard single-case methods and group-level reference data.

### 3. fMRI localizers

#### The language localizer

This language localizer task, introduced in Fedorenko et al. (9) and used in many subsequent studies (e.g., (10–15); the task is available for download from https://www.evlab.mit.edu/re-sources) included two conditions in a blocked design. Participants silently read sentences and lists of unconnected, pronounceable nonwords in a blocked design (**Fig. 3B**). The Sentences > Nonwords contrast targets cognitive processes related to high-level language comprehension, including understanding word meanings and combinatorial linguistic processing. Each stimulus (sentence or nonword list) was 6 seconds long and consisted of 12 words or nonwords presented one word/nonword at a time at the rate of 450 ms per word/nonword. The main task was attentive reading. Each stimulus was followed by a simple button-press task, which was included to maintain alertness. Trials were grouped into blocks of 3 trials of the same condition (18 seconds total). Each scanning run consisted of 16 blocks (8 per condition) and 5 blocks of a baseline blank screen (14s each), for a total run duration of 358 seconds. Condition order was counterbalanced across runs. Each participant completed 2 runs.

#### The Multiple Demand system localizer

This spatial working memory localizer task, introduced in Fedorenko et al. (16) and used in many subsequent studies as a localizer for the MD system (17–23), included two conditions in a blocked design. Participants had to keep track of spatial locations presented in a sequence (8 locations in the Hard condition, 4 locations in the Easy condition) (**Fig. 2E**). The Hard > Easy contrast targets cognitive processes broadly related to performing demanding tasks—what is often referred to by an umbrella term ‘executive function processes’. Each trial consisted of a brief fixation cross shown for 500 ms followed by 4 sequential flashes of unique locations within the 3 × 4 grid (1 s per flash; two locations at a time in the Hard condition, one location at a time in the Easy condition). Each trial ended with a two-alternative, forced-choice question (two sets of locations were presented for up to 3.25 s, and participants had to choose the set of locations they just saw; if they responded before 3.25 s elapsed, there was a blank screen for the remainder of the 3.25 s period). Finally, participants were given feedback in the form of a green checkmark (correct response) or a red cross (incorrect response or no response) shown for 250 ms. The total trial duration was 8 seconds. Trials were grouped into blocks of 4 trials of the same condition (32 seconds total). Each scanning run consisted of 12 blocks (6 per condition) and 4 blocks of a baseline fixation screen (16 seconds each), for a total run duration of 448 seconds. Condition order was counterbalanced across runs. Each participant completed 2 runs.

### 4. fMRI acquisition, preprocessing, modeling, and analyses

#### fMRI data acquisition

Whole-brain structural and functional data were collected on a whole-body 3 Tesla Siemens Trio scanner with a 32-channel head coil at the Athinoula A. Martinos Imaging Center at the McGovern Institute for Brain Research at MIT. T1-weighted structural images were collected in 176 axial slices with 1 mm isotropic voxels (repetition time (TR) = 2,530 ms; echo time (TE) = 3.48 ms). Functional, BOLD data were acquired using an EPI sequence with a 90° flip angle and using GRAPPA with an acceleration factor of 2; the following parameters were used: 31 4.4 mm thick near axial slices acquired in an interleaved order (with 10% distance factor), with an in-plane resolution of 2.1 × 2.1 mm, FoV in the phase encoding (A >> P) direction 200 mm and matrix size 96 × 96 voxels, TR = 2,000 ms and TE = 30 ms. The first 10 s of each run were excluded to allow for steady state magnetization.

#### fMRI data preprocessing

fMRI data were preprocessed and analyzed using SPM12 (release 7487), CONN EvLab module (release 19b), and other custom MATLAB scripts. Each participant’s functional and structural data were converted from DICOM to NIFTI format. All functional scans were coregistered and resampled using B-spline interpolation to the first scan of the first session (24). Potential outlier scans were identified from the resulting subject-motion estimates as well as from BOLD signal indicators using default thresholds in CONN preprocessing pipeline (5 standard deviations above the mean in global BOLD signal change, or framewise displacement values above 0.9 mm; (25)). Functional and structural data were independently normalized into a common space (the Montreal Neurological Institute [MNI] template; IXI549Space) using SPM12 unified segmentation and normalization procedure (26) with a reference functional image computed as the mean functional data after realignment across all timepoints omitting outlier scans. The output data were resampled to a common bounding box between MNI coordinates (−90, −126, −72) and (90, 90, 108), using 2 mm isotropic voxels and 4th order spline interpolation for the functional data, and 1 mm isotropic voxels and trilinear interpolation for the structural data. Last, the functional data were smoothed spatially using spatial convolution with a 4 mm FWHM Gaussian kernel.

#### fMRI data modeling

For all experiments, effects were estimated using a general linear model (GLM) in which each experimental condition was modeled with a boxcar function convolved with the canonical hemodynamic response function (HRF) (fixation was modeled implicitly, such that all timepoints that did not correspond to one of the conditions were assumed to correspond to a fixation period). Temporal autocorrelations in the BOLD signal timeseries were accounted for by a combination of high-pass filtering with a 128 s cutoff, and whitening using an AR (0.2) model (first-order autoregressive model linearized around the coefficient a = 0.2) to approximate the observed covariance of the functional data in the context of restricted maximum likelihood estimation. In addition to experimental condition effects, the GLM design included first-order temporal derivatives for each condition (included to model variability in the HRF delays), as well as nuisance regressors to control for the effect of slow linear drifts, subject-motion parameters, and potential outlier scans on the BOLD signal.

#### fROI definition and response estimation

Definition of the language fROIs: Each individual map for the Sentences > Nonwords contrast from the language localizer was intersected with a set of 5 parcels. These parcels (available at https://www.evlab.mit.edu/resources) were derived from a probabilistic activation overlap map for the same contrast in a large set of independent participants (n=220) and covered the frontotemporal language system in the left hemisphere (27). Within each parcel, a participant-specific language fROI was defined as the top 10% of voxels with the highest t-values for the localizer contrast. To estimate the response in the language fROIs to the conditions of the language localizer, the same cross-validation procedure was used as described above, to ensure independence. As expected, the language fROIs showed a robust Sentences > Nonwords effect (ps < 0.001, |d|s > 2.05). Similarly, for the definition of the MD fROIs: Each individual map for the Hard > Easy spatial working memory contrast from the MD localizer was intersected with a set of 20 parcels (10 in each hemisphere). These parcels (available at https://www.evlab.mit.edu/resources) were derived from a probabilistic activation overlap map for the same contrast in a large set of independent participants (n=197) and covered the frontal and parietal components of the MD system bilaterally (28, 29). Within each parcel, a participant-specific MD fROI was defined as the top 10% of voxels with the highest t-values for the localizer contrast. To estimate the response in the MD fROIs to the conditions of the MD localizer, the same cross-validation procedure was used as described above. As expected, the MD fROIs showed a robust Hard > Easy spatial working memory effect (ps < 0.001, |d|s > 1.84; here and elsewhere, p-values are corrected for the number of fROIs using the false discovery rate (FDR) correction (30)).

#### Statistical models

Linear mixed-effects models were implemented in R (lme4 package; (31)), with random intercepts for participants and regions. P-values were approximated using the lmerTest package (32), and effect sizes (Cohen’s *d*) were estimated using the EMAtools package (33).

#### A whole-brain GSS analysis

To search for areas sensitive to deductive reasoning load, we used a group-constrained subject-specific (GSS) analysis (9, 34), which is similar to a random-effects group analysis but allows for inter-individual variability in the precise locations of functional areas. This analysis identified a few areas in the frontal and parietal cortex that responded strongly to the deduction contrast (in left-out data). The responses in the MD and deductive-reasoning fROIs were statistically evaluated similarly to the language fROIs’ responses.

